# Global success in oyster reef restoration despite ongoing recovery debt

**DOI:** 10.1101/2022.01.23.477429

**Authors:** Deevesh A. Hemraj, Melanie J. Bishop, Boze Hancock, Jay J. Minuti, Ruth H. Thurstan, Philine S.E. Zu Ermgassen, Bayden D. Russell

## Abstract

Habitat destruction and biodiversity loss from exploitation of ecosystems have led to increased restoration and conservation efforts worldwide. Disturbed ecosystems accumulate a recovery debt – the accumulated loss of ecosystem services - and quantifying this debt presents a valuable tool to develop better ecosystem restoration practices. Here, we quantified the ongoing recovery debt following structural restoration of oyster habitats, one of the most degraded marine ecosystems worldwide. We found that whilst restoration initiates a rapid increase in biodiversity and abundance of 2- to 5-fold relative to unrestored habitat, recovery rate decreases substantially within a few years post-restoration and accumulated global recovery debt persists at >35% per annum. Therefore, while efficient restoration methods will produce enhanced recovery success and minimise recovery debt, potential future coastal development should be weighed up against not just the instantaneous damage to ecosystem functions and services but also the potential for generational loss of services and long-term recovery.

## Introduction

Exploitation and disturbance of ecosystems in the Anthropocene has led to severe degradation of natural biomes and loss of biodiversity^1,2,3^. Consequently, investment in conservation and restoration efforts have increased worldwide^4,5,6,7^, especially as a strategy to restore ecosystem services^8^. Whilst the cost-benefit ratio of restoration is often justified as ecosystem recovery that yields sufficient benefits to human prosperity^9^, recovery of ecosystems back to a reference state in terms of biodiversity and ecosystem functions and services^10^ often requires decades^11,12,13^. Where damaged ecosystems provide reduced function or support reduced biodiversity relative to the historical “natural state” (reference/pristine condition), a recovery debt is accumulated^13^ (Fig. 1). While recovery debt has been estimated in ecosystems that largely only require natural regeneration following the removal of persistent disturbances^13^, the recovery debt and recovery pathway of marine habitats requiring active intervention, including structural restoration, remains undetermined (Fig. 1).

**Fig. 1:**
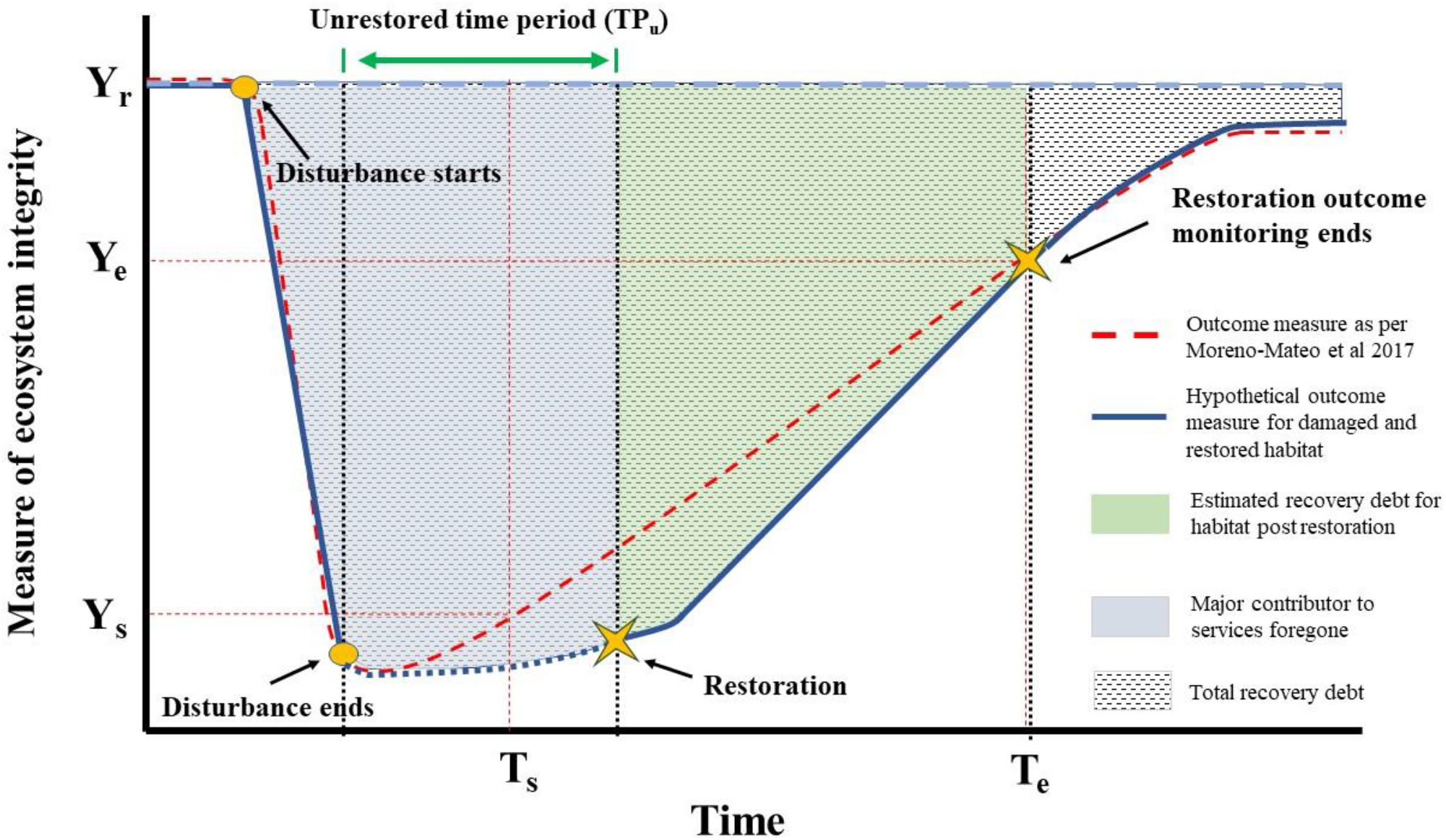
Theoretical diagram of general recovery debt (red dotted line) and recovery debt specific to restored habitats (blue lines). TP_u_ reflects the ecosystem integrity in the absence of restoration efforts Y_s_ and T_s_ represent ecosystem integrity outcome measure and time when measurement started. Y_e_ and T_e_ represent ecosystem integrity outcome measure and time when measurement ended. Pale blue dotted line (Y_r_) represents ecosystem integrity outcome measure of reference site. Note that Time (x-axis) is not to scale and the unrestored time period (TP_u_) from when disturbance stopped to restoration could be 20 - 50-fold longer than the post-restoration period; in some cases, TP_u_ can be over 100 years. Figure modified from Moreno-Mateos et al. (2017).

A major part of the accumulated debt in recovering ecosystems can be considered as services foregone^14^ (future services had there not been damage). Actions which increase the rate of system recovery (e.g., habitat restoration) will theoretically increase both the rate of recovery and the potential for an ecosystem to recover to its maximum capacity, minimise the services foregone, and thus reduce recovery debt. Therefore, utilising the best performing restoration methods to rapidly boost recovery of ecosystems may minimize the accumulated recovery debt and at least partially offset the ongoing damage associated with current activities (e.g., coastal development).

Oyster habitats are one of the most anthropogenically impacted coastal habitats worldwide. At least 85% of oyster habitats have been lost globally, predominantly as a consequence of historical overharvest using destructive fishing practices, but also due to more recent effects of coastal urbanisation, including declining water quality and introduced disease issues^15,16^. Destructive dredge harvest not only removed live oysters and their biological functions, but also the remnant dead oyster shells that provide structural complexity (vertical relief and size) and substrate for oyster settlement^15^. As such, only a handful of sites remain globally where oyster habitats remain in their ‘natural state’ (mainly in the East and Gulf Coasts of the USA). Given the biogenic reef building nature of oyster habitats, and a life history that leaves them vulnerable to allee effects, natural recovery is unlikely given the loss of structural habitat which is essential for oyster settlement. Therefore, intensive restoration efforts of oyster habitats have led to large capital investment in various methods, all aiming to increase the spatial area of oyster habitat, their functioning, and ecosystem services^17,18,19,20,21,22^.

Restoration of oyster habitats typically includes remediation of environmental conditions, substrate provision and/or restocking with juvenile and/or adult oysters^23^. Key considerations in substrate provision are the type of material used (e.g., recycled shell vs. artificial materials such as concrete blocks) as well as its spatial arrangement^24^. While oyster habitats naturally accrete on oyster shell, the availability of oyster shells (from aquaculture, or shell recycling program) is generally limiting, meaning that different substrate types have been tested as an alternative in oyster habitat restoration studies. Although a range of factors associated with the spatial arrangement of substrate (e.g., patch size, fragmentation) can influence oyster establishment and ecosystem service provision^24^, vertical relief is considered particularly important as it can influence oyster habitat growth by determining water flow, dissolved oxygen concentrations, and reduce smothering from the accumulation of sediment.

The exploitation and removal of oyster habitats largely took place during, or prior to, the 19^th^ century^25,26,27^. While scarce documentation exists which depicts the pristine or pre-impact condition of oyster reefs, it is widely accepted that our current understanding is hampered by a shifted baseline. Recovery debt following structural restoration of oyster habitats (e.g., Fig. 1) can therefore currently only be assessed relative to remnant habitat (Box 1). Assessment of the current recovery can be used to identify the extent to which restoration efforts can mitigate contemporary damage (e.g., with coastal development), to improve the incorporation of recovery debt in restoration planning, environmental offsets, and in mitigation measures. While oyster habitat restoration tends to yield some positive results in terms of recovery towards a reference state, the effectiveness of varying methods of restoration in terms of maximum habitat recovery remain unclear. Here, we calculated the recovery debt for restored oyster habitat globally and undertook a meta-analysis of oyster habitat restoration worldwide to: (1) calculate restoration associated recovery of biodiversity and abundance of resident and transient fish and invertebrates in oyster habitats; and (2) identify the methods for oyster habitat restoration which most successfully reduced recovery debt. Overall, we demonstrate that restoration is effective at rapidly mitigating damage to oyster habitat ecosystems, but while the accumulated debt is variable among different measures of recovery, debt continues to accumulate.

### Box 1: Key Terminologies

**Oyster habitat**: a patch of oysters large enough to form three-dimensional complex habitat. Similar terminology used in the literature include ‘oyster bed’ or ‘oyster reef.

**Recovery debt**: accumulated loss of ecosystem structure and functions between the point of habitat damage and “full recovery” to a reference state.

**Restored habitat**: an oyster habitat patch that has been actively restored, for example, by the addition of substrate (e.g., oyster shell, limestone, concrete) and/or the provision of live oysters

**Remnant habitat**: oyster habitats that have not been destroyed or degraded (e.g., by extraction of oysters) and have persisted over centuries, or those that have historically been damaged but have since fully recovered through natural processes. These habitats are used as reference habitat for calculating recovery debt of restored reefs.

**Unrestored habitat**: an area where oysters historically were present but are presently degraded and are not being restored. These habitats are generally areas of bare sediment where oyster reefs previously existed.

## Results

### Oyster habitat recovery post restoration

The analysis of monitoring data for 20 restored oyster habitats, obtained over an average of four years post restoration (Fig. 2a, b), revealed that the restored habitats had an annual average of 36.08% (± 5.58 SE) lower species diversity of fish and invertebrates than remnant habitats. While four restoration sites recovered well in terms of diversity within three to four years post restoration (RDr < 10%), all remaining sites had a recovery debt of >20% (Fig. 2). Total abundance of fish and invertebrates recovered better than diversity, having a mean recovery debt of 24.37% per annum (± 9.28 SE), over an average monitoring period of 3 years. In contrast to diversity, fish, and invertebrate abundance at 5 out of 20 restored habitats had fully offset the recovery debt (negative recovery debt) after two and a half years, suggesting complete recovery and even higher fish and invertebrate abundance compared to remnant habitats. It must be noted, however, that abundance does not account for shifts in relative abundance among species compared to remnant habitats and does not discriminate between attraction and production.

**Fig. 2.**
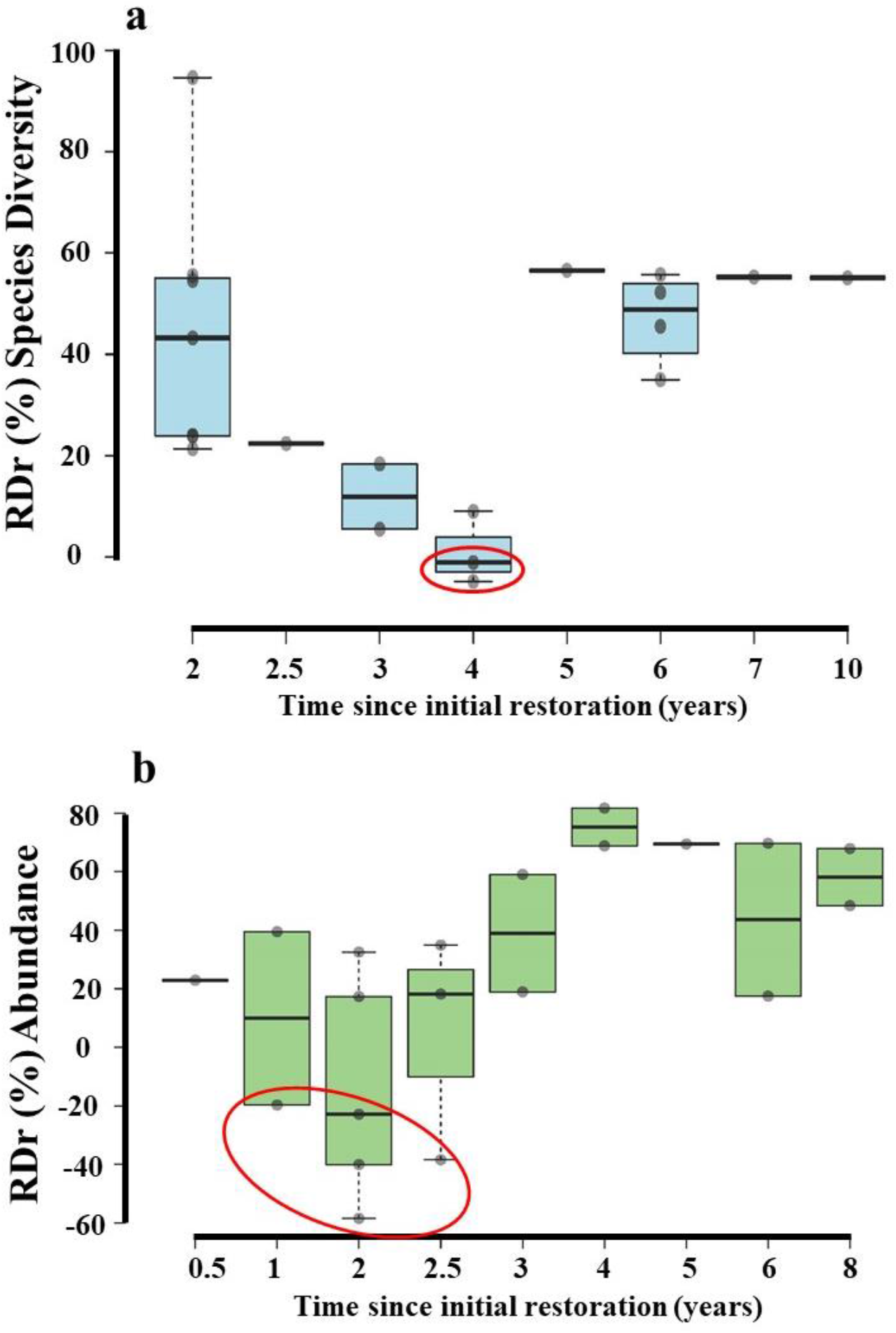
Oyster habitat accumulated recovery debt per annum as a function of time since restoration. **a-b** Recovery debt calculated from diversity (a) and abundance (b) (n = 20 sites for each). Accumulated debt declines initially with rapid recovery following restoration, but then begins to increase as recovery slows and debt begins to accumulate again. Black dots represent estimated recovery debt data points. Black lines represent median recovery debt. Box limits represent 25^th^ and 75^th^ percentile. Note the different scales of each graph. Red circles represent data points extracted from studies that used limestone, oyster shell or a combination of both as substrate for habitat building^28,29,30^.

Over the longer-term, neither diversity nor abundance showed a consistent relationship between estimated recovery debt and time (years) since implementation of structural restoration (Fig. 2; r^2^ = 0.029, P = 0.458, r^2^ = 0.057, P = 0.315, respectively). However, during the first 4 years, there was substantial decrease in recovery debt in terms of species diversity (slope: −22.849, r^2^: 0.4962, P = 0.0054). Annual recovery rates were high in the first two to four years but then decreased (Fig. 3). Overall, with a few exceptions, restored oyster habitats tended to recover towards a reference state (Fig. 3; percentage recovery rate = 27.05 ± 4.07 SE and 90.16 ± 32.16 SE for diversity and abundance, respectively) though there was no indication as to when, or if, the habitats would reach “full recovery” (matching reference habitats).

**Fig. 3.**
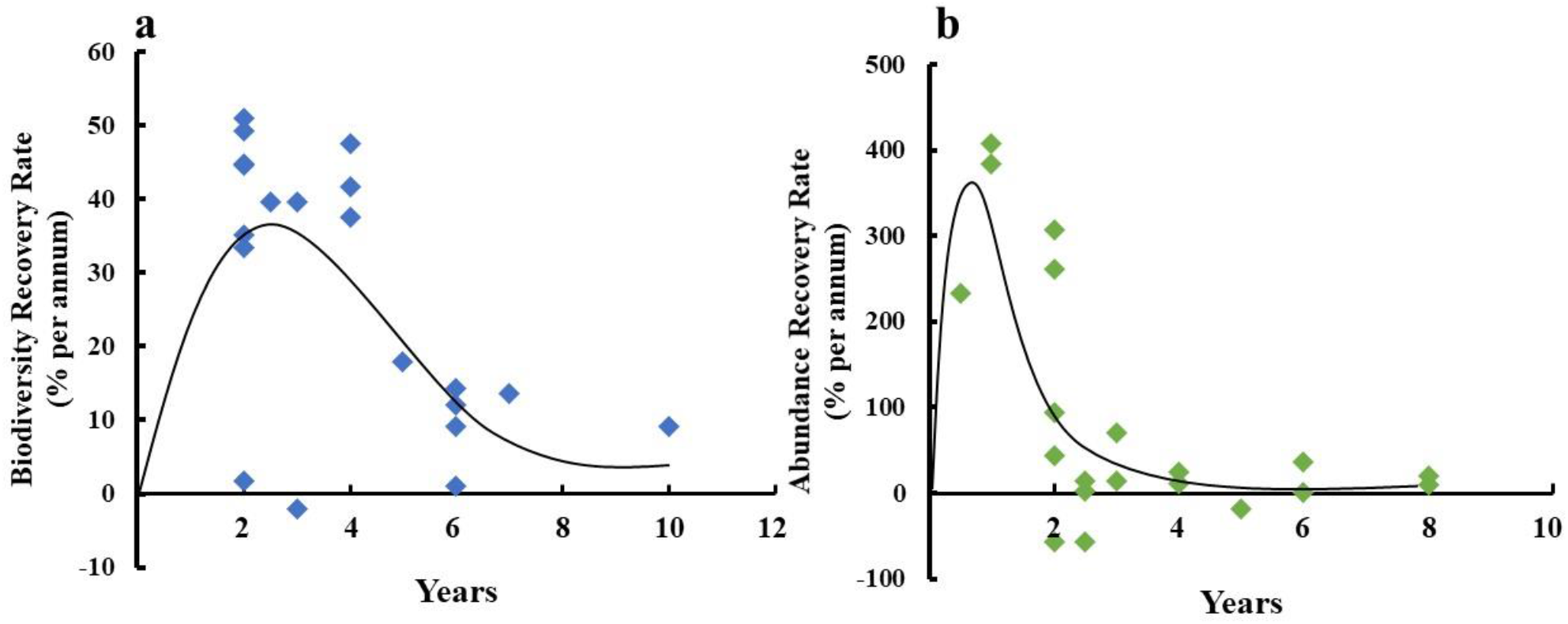
Oyster habitat recovery rates against time of monitoring. **a-b** calculated recovery rate using fish and invertebrate diversity (a) and abundance (b). Note the different scales for diversity and abundance indicating that abundance has much more rapid recovery than diversity. The black lines represent a smoothed quadratic model with intercept set at 0. Recovery rates are calculated in relation to a reference remnant site.

### Difference in diversity and abundance between restored and unrestored habitats

The calculated effect sizes (lnRR) indicated that compared to unrestored habitats (areas that have been left in a degraded state for decades (generally as bare sediment), restored habitats had an overall greater nekton abundance, (δ = 1.117 ± 0.309, P < 0.001) (see Supplementary File 2 for all meta-analysis results). Invertebrate abundance displayed a larger effect size between restored and unrestored habitats than fish abundance (93.5% increase for fish and 532.2% increase for invertebrates) though both were significant (invertebrates: δ = 0.273 ± 0.264, P < 0.042; fish δ = 1.294 ± 0.48, P < 0.001; Fig. 4a, b). The effect size for abundance was greatest in the first year of habitat restoration, and overall displayed a negative relationship with time (Q = 7.76, df = 1, P = 0.005; Fig. 4c), suggesting that following a period of rapid response, recovery slowed. Yet, while the rate of increase in abundance declined over time (Fig. 3), abundance remained consistently higher in restored habitats relative to unrestored sites (Fig. 4c).

**Fig. 4.**
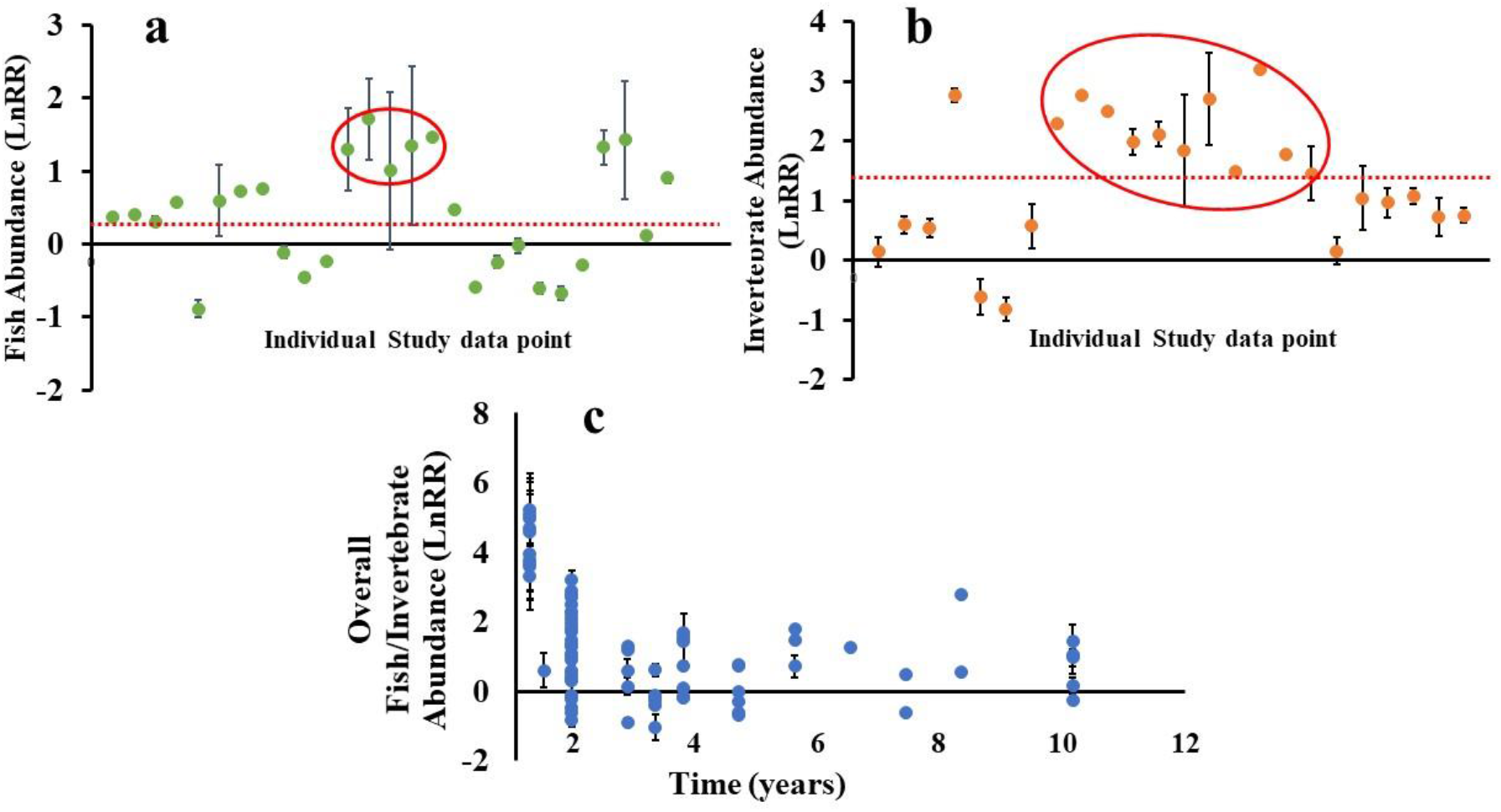
Inverted forest plots representing effect size for increase or decrease in transient and resident fish and invertebrate relative to unrestored habitats. **a-b** Change in fish (a) and invertebrate (b) abundance in restored oyster habitats compared to bare sediment. Data points = effect sizes (lnRR). X-axes in graphs (a) and (b) only represents distribution of data points. Red dotted line represents overall mean effect size. Red circles represent data points extracted from studies that used limestone, oyster shell or both as substrate for habitat building^29,31,49^. **c** Overall abundance of oyster habitat associated fauna remains higher than that of bare sediment over time. Error bars = 95% CI. Data points without visible error bars are due to very small CI.

### Oyster habitat restoration method

Overall, more oyster spat recruited to oyster shells than 15 alternate substrata (of which limestone, concrete, and granite were most common) (δ = −0.472 ± 0.203, P < 0.001; Fig. 5a). Of the alternate substrata, limestone performed the closest to oyster shells with no significant difference between the two (δ = 0.120 ± 0.256, P = 0.356; Fig. 5a). Granite seemed to attract slightly fewer recruits than oyster shells (7 of 12 studies), but that difference was not significant (δ = −0.206 ± 0.657, P = 0.540). However, fewer recruits (approximately −37% compared to oyster shell) settled on concrete structures (δ = −0.788 ± 0.372, P < 0.001; Fig. 5a). Restored habitats which recouped the recovery debt (negative debt) and had the greatest increase in abundance^28,29,30,31^ were all constructed with either limestone, oyster shell, or a mix of both (Fig. 2, 4).

**Fig. 5.**
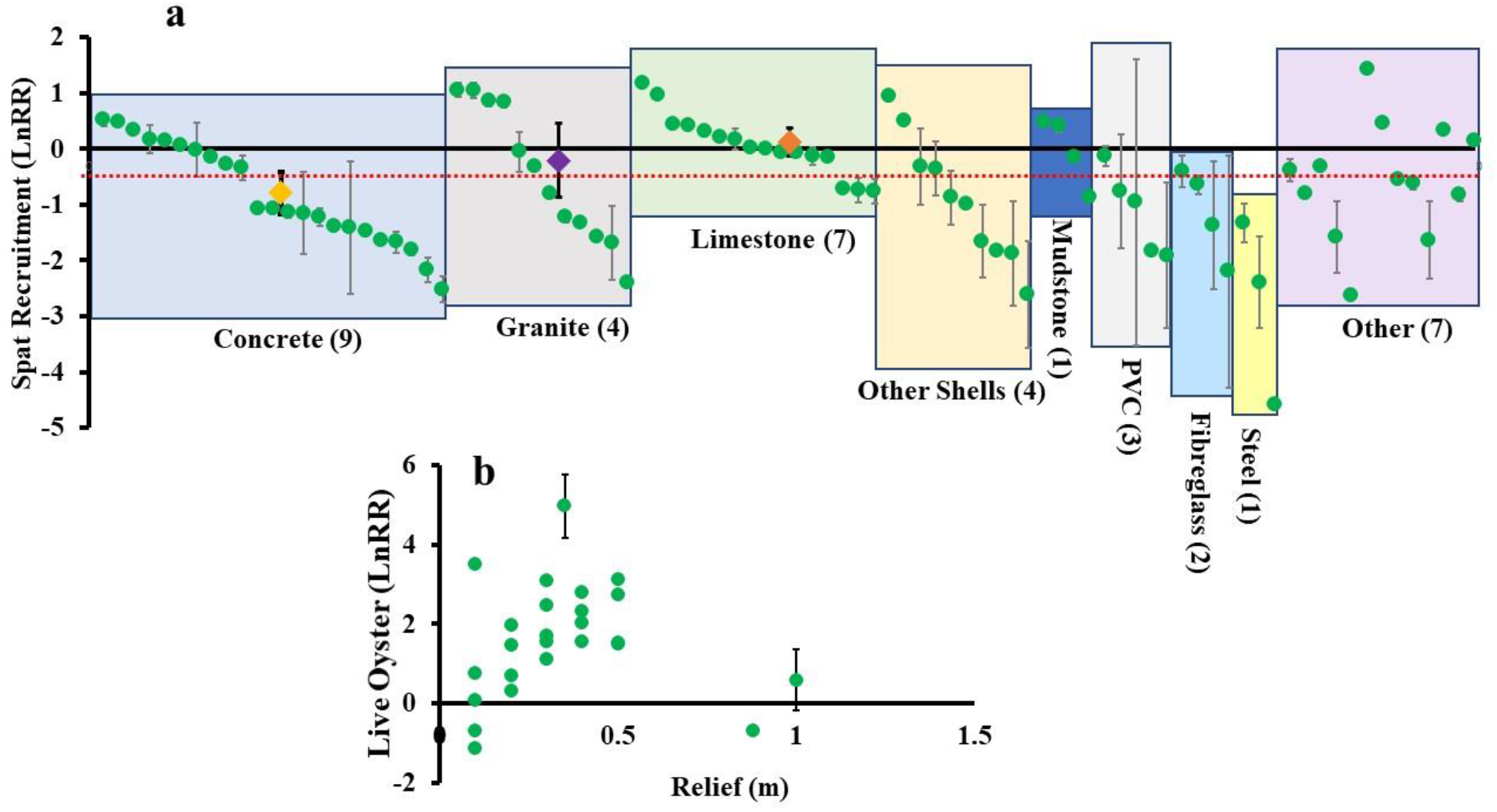
Inverted forest plot representing difference in overall spat settlement and oyster density. **a-b** Spat settlement on alternative substrata compared to oyster shell (a) and change in live oyster density on oyster habitats as a function of vertical relief above the sediment (b). Data points = effect size (lnRR). Error bars = 95% CI. Data points without visible error bars are due to very small CI. Yellow, purple, and orange diamonds represent the mean effect sizes for concrete, granite, and limestone, respectively. Red dotted line represents overall mean effect size for all alternative substrata compared to using oyster shells. Numbers in parentheses represent the number of papers from which the data points were taken.

Vertical relief influenced the density of live oysters whereby oyster habitats more than 20 cm above the sediment had ~84% higher live oyster density than unrestored bare sediment (δ = 1.771 ± 0.474, P < 0.001; Fig. 5b) while oyster habitats with vertical relief <20 cm did not support higher oyster densities than unrestored bare sediment (δ = 0.34 ± 1.391, P < 0.631). No linear relationship was found between relief and oyster density, with increased vertical relief above 20 cm not contributing to substantially more recruitment (Q = 0.0715, df = 1, P = 0.789, Fig. 5b).

## Discussion

Historical exploitation has left the majority of ecosystems formed by oysters in a severely degraded state for decades to centuries. Our analyses focus on the contemporary debt that is still accrued following restoration meaning that mitigation of the damage to coastal habitats will, at the very least, take more direct intervention and time to recover than generally anticipated. We found that recovery debt tends to decrease during the immediate 2 - 4 years following restoration across all the locations assessed globally, concomitant with rapid colonisation of biota - an important result given the increasing investment of resources in oyster habitat restoration worldwide. The decrease in recovery debt is not, however, maintained through time and following a rapid initial recovery of faunal assemblages associated with the restored habitat, reducing the accrued debt, there is a gradual increase in debt as recovery slows (Fig. 2 and 3). This shift likely reflects an initial rapid accumulation of biodiversity of early successional species, followed by establishment of competitively dominant taxa that stabilize the assemblage structure and exclude some species. This initial increase in species abundance/diversity followed by subsequent community turnover and change in species interactions is a trend of recovery through time observed in many terrestrial and aquatic ecosystems^32,33,34^. In fact, ecosystem complexity and recovery are attained following build-up of species abundance and richness, community turnover, and meta-community interactions^11,12,13^. Therefore, while restoration can be effective in rapidly reducing the debt that accrues following destruction of coastal habitats, focusing monitoring on the initial years following restoration will overestimate the trajectory towards recovery ^35^.

The recovery of diversity of oyster habitat-associated fish and invertebrates was slower than that of abundance (~36% and ~24% recovery debt, respectively). This differs from the previous estimates from most ecosystems whereby overall recovery debt in diversity is generally higher than that of abundance^13^. The trajectory of recovery in abundance and diversity tend to differ in ecosystems depending on the type of restoration practice (active vs. passive restoration) by either driving rapid abundance of opportunistic colonisers or slow progression in community turnover^36^. For example, in the terrestrial realm, active landscape restoration (e.g., tree planting) tends to increase faunal abundance faster than diversity because of the sudden change in habitat structure which can be rapidly exploited by few species (e.g., forest specialists^34,36^). On the other hand, similar barren landscapes undergoing passive recovery will experience progressive community turnover from an open-field community to a forest species dominated community as the habitat setting gradually changes^36^. Comparable trends have been recorded in active mangrove restoration whereby abundance of algivorous fish species peak after restoration, but overall fish diversity remains low^37^. While the progression in recovery of abundance and diversity of organisms have not been contrasted between passive or active restoration efforts in multiple marine habitats, our results suggest a fast increase of abundance of some species in restored oyster habitats, where active restoration by substrate provision is generally unavoidable.

It is likely that attraction of mobile fauna from adjacent habitats to the more structurally complex restored habitats, rather than purely enhanced recruitment, accounts for some of this rapid increase in faunal abundance^38,39^. Interestingly, 25% of restored sites we assessed gained higher abundance than their reference sites (remnant habitats). As the remnant habitats themselves are likely to have experienced some extent of change since industrial overfishing began in the 19^th^ century (shifting baselines), this higher abundance is likely to be, at least partly, reflective of somewhat disturbed remnant habitats. Unfortunately, the multigenerational exploitation and damage of marine systems means that we have lost most undisturbed “reference” baselines. Anecdotally, many of the ‘remnant habitats’ are actually reefs formed from other human activities like abandoned benthic oyster farm infrastructure or even discarded rock ballast from early trade, making it largely impossible to quantify the degree of this past impact; effectively we cannot recreate the true historical baseline. Many of our estimates consider locations where nominally undisturbed remnant habitats were available for comparison with restored habitats, yet it is important to note that these locations form a very small proportion globally of the areas where habitats would have been historically^15^. In addition, the short duration of most monitoring programmes (2-6 years) means that it is not possible to quantify the time to full recovery. Nonetheless, our estimated recovery debt, along with the considerable decrease in the rate of recovery over time, suggest that an initial rapid partial recovery of oyster habitat associated fish and invertebrates is likely in restored habitats, but complete recovery for both abundance and diversity will require >10 years (Fig. 6).

**Fig. 6.**
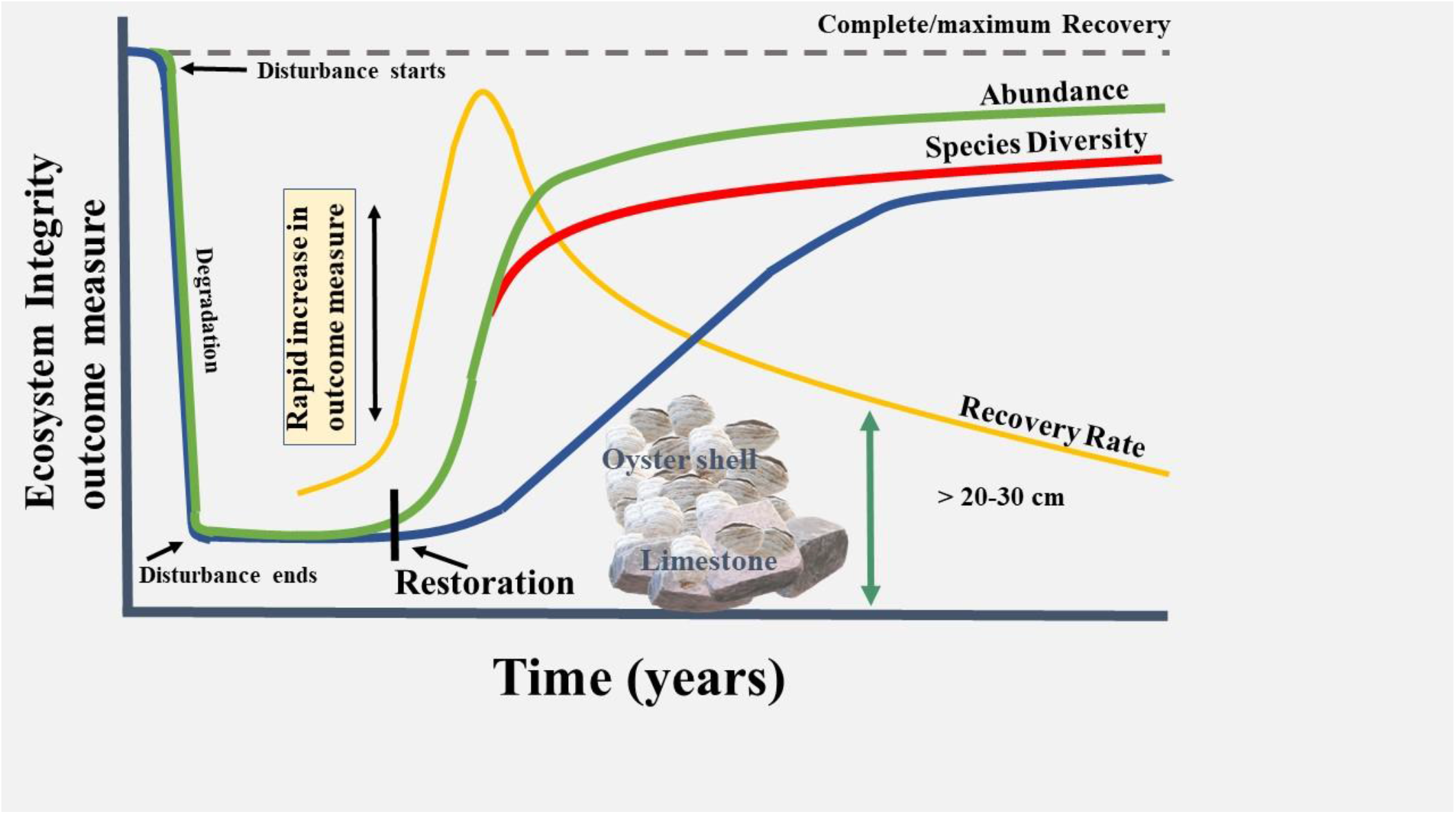
Model of oyster habitat recovery following disturbance and subsequent restoration. Trends are based on analysis of change in overall recovery of oyster habitat (blue line), cumulative species diversity (red line) and cumulative abundance (green line) of associated species. Note the initial rapid recovery rates post-restoration (yellow line) which then declines over time.

Irrespective of the accrued recovery debt, restoration efforts rapidly increase habitat function relative to unrestored sites. Restoration contributes to approximately double the abundance of fish and more than fivefold the abundance of invertebrates to coastal ecosystems over unrestored habitats. Such increases are promising in terms of recouping ecosystem services such as fisheries^22,39,40^. For example, multiple assessments grounded on the increase in habitat provisioning and nekton abundances show that restoration provides multiple prospects for fisheries^22,41,42,43^. Nonetheless, the general temporal progression of ecosystem recovery towards climax community composition through compositional turnover^44^, community/meta-community interactions, and broader ecosystem resilience and stability have to be accounted for when managing ecosystem recovery^2,12,41^. In this sense, complementing active restoration with adequate time and protection for the habitat to mature will further benefit recovery^42^.

While oyster habitat restoration is generally beneficial in terms of increased oyster density and oyster habitat associated biodiversity, not all restoration methods performed equally. First, we found that oyster shell was the best substrate for habitat building in terms of spat recruitment. However, oyster shells are not readily available in bulk for large scale restoration, may have biosecurity risks if not adequately weathered prior to use, may not provide sufficiently stable structure in wave-swept areas, and have high monetary costs^43^. We also advise caution with the use of other types of shells, as there is preliminary evidence that brittle or thin shells may break down rapidly and not form the structure which is key for spat recruitment and survival (e.g., the use of surf clam shell in Harris Creek, Chesapeake Bay^44^). As an alternative substrate when oyster shell is limited, limestone performed almost as well in terms of spat recruitment. In fact, the best performing restoration projects from our analysis (e.g., *C. virginica* reefs in the USA) used either oyster shell, limestone, or a mix of both as substrate for habitat building^28,29,30,31,49^. Secondly, our finding that live oyster density is maximised on habitats with structure more than 20-30 cm above the sediment reinforces current restoration practices^50,51,52^. Habitats with higher relief are more likely to avoid smothering of oysters by sedimentation and elevate oysters above seasonally hypoxic bottom waters thereby increasing survival of spat and adults^51,53^. The maximum relief of habitats above the sediment will be defined by water depth and tidal range, especially for intertidal habitats. Such intertidal habitats will expand laterally, gaining surface area rather than height, while subtidal habitats have the potential for both lateral and vertical growth. Irrespective of whether restoration is inter- or subtidal, however, we demonstrate that greatest success is achieved when the restoration substrate is sufficiently above the sediment, providing refined guidance for restoration planning.

Overall, we demonstrate that active restoration of oyster habitats provides enormous benefits to the recovery of associated faunal diversity and abundance (Fig.4). Our measurement of recovery debt post-restoration highlights that recovery of degraded oyster habitats to a reference state is a long-term process and will also benefit from elimination of any external disturbance (e.g., protection from oyster harvest). In addition, ecosystems require time to develop a stable and resilient community structure following active structural restoration. Nonetheless, implementing the appropriate restoration methods has the potential to boost recovery rate, improve overall outcomes, and maximise return for effort. It must be noted that, currently, monitoring of restored habitats is generally done for < 5 years post-restoration, capturing the initial boost in recovery but not the subsequent progressive change in community composition that remains integral to regaining full ecosystem complexity^12^. Refining our understanding of the capacity of restored habitats to recover full functions and services will require longer-term monitoring, even more so in areas where remnant reference reefs are not present as maximum recovery in such habitats will only likely be indicated by long-term maintenance of ecosystem complexity and stability. From a different perspective, we bring into focus that the actions to offset or mitigate the damage caused by coastal development may be inadequate and the prospect of future sustainable development should be weighed up against not just the instantaneous loss of ecosystem function and services, but the potential for generational loss as has been the case for oyster habitats. Overall, by integrating an estimation of oyster habitat recovery with an assessment of the most effective restoration methods we show that, globally, biodiversity and abundance benefit immensely from oyster habitat restoration and the recovery completeness will progressively increase on potentially decadal scales.

## Methods

### Literature search

Our analysis followed the PRISMA (Preferred Reporting Items for Systematic Reviews and Meta-Analyses) and the CEE (Collaboration for Environmental Evidence) guidelines. We aggregated studies targeting oyster habitat restoration by using the search terms ((“oyster reef” OR “oyster habitat” OR “oyster bed”) AND (“restoration” OR “recovery” OR “rehabilitation” OR “substrate” OR “relief” OR “biodiversity” OR “species richness” OR “abundance” OR “living shoreline” OR “community” OR “epifauna” OR “nekton”)) from three databases: Google Scholar, Scopus and Web of Science. Study identification was terminated on the 29^th^ of September 2021 (range: 1970 to 29^th^ September 2021) and only peer-reviewed journal articles and dissertations were included in our study. Also, we used species abundance and diversity for recovery debt and rate calculations as few papers documented how other parameters (e.g. filtration, wave attenuation) changed post restoration compared to a remnant site (low sample size). Our initial literature search yielded 12,128 papers. After removal of duplicates and studies that were out of context, 1,374 papers remained (Primary screening; Supplementary File 1). We then screened these papers to identify those that were specifically relevant to oyster restoration projects. The majority of studies (~73%) and sites focusing on oyster habitat restoration were situated in the east coast of North America (Fig. S1 and S2).

### Selection criteria

We removed duplicate papers and manually screened the titles and abstracts of each study to select studies that explicitly targeted oyster habitat restoration. We included all papers that studied one or more of the following:

1. A measure of the resident or transient fish and invertebrates sampled in restored and remnant habitats (e.g., abundance, density, CPUE, species richness, diversity).
2. A measure of the resident or transient fish and invertebrates sampled in restored oyster habitats and degraded habitat (commonly represented as bare sediment).
3. A measure of oyster density in relation to oyster habitat vertical relief
4. A measure of recruitment on oyster shell and other substrata for restoration.

To be extracted and used in our analysis, studies had to report data either as mean/median with a measure of variance (e.g., SD or range) in tables or figures, or provide the full data set from which mean, and SD could be calculated. In the case a study reported data from multiple sites, each site was used as an individual data point. If a study reported two metrics that were of interest (e.g., diversity and abundance, or fish abundance and invertebrate abundance), each metric was analysed separately and as appropriate for our analysis. We only included data which were directly relevant to oyster habitat performance, excluding anything that could indirectly come from the influence of other types of habitats (e.g., adjacent marsh or mangroves). For example, if a study reported a metric from a control site, an oyster-only site and an oyster and seagrass site, we only use the data from the control and oyster-only sites. When studies reported data over shorter time intervals than yearly (e.g., monthly), we calculated a pooled annual mean and SD including each data point in our estimation to capture the whole range of response^54^. Based on the selection criteria for our research question, data were then extracted from 70 papers spanning sites worldwide (Supplementary Fig. 1). From these papers, a total of 232 data points were retrieved to estimate recovery debt in terms of biological diversity (n = 20 data points) and transient and resident fish and invertebrate abundance (n = 20), to analyse difference in fish and invertebrate abundance between restored and unrestored habitats (n = 76), estimate the influence of different substrates on oyster spat recruitment (n = 90), and estimate the influence of vertical relief on oyster density (n = 26). Data for analysis were extracted from figures using PlotDigitizer for windows, or from tables and text.

### Calculating recovery debt and recovery rate

Recovery debt was calculated following^13^. In brief, we screened all studies that reported an outcome metric that was either species richness, diversity index, species density, or species abundances. Here we used overall organism diversity or abundance (combining fish and invertebrates) linked to reef restoration to obtain the best estimate of overall recovery debt for each reef. For recovery rate and debt calculations we only used data from studies that included the outcome metrics (e.g., abundance and diversity metrics) from before restoration and after restoration (no matter the time post restoration), at the restoration and a reference remnant site. Recovery debt in terms of diversity (including metrics representing the number of species utilising a site, e.g., species richness and diversity) and abundance (including metrics representing an estimate of the number of individuals within a site, e.g., abundances, CPUE and density) were then separately calculated using the following equations:

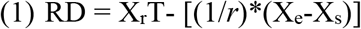

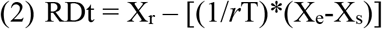

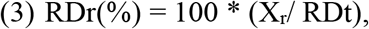

where, RD is the estimated graphical area of recovery debt (Fig. 1) for the time period where monitoring took place, RDt is the of recovery debt per annum, and RDr(%) is the estimated percentage recovery debt per annum. X_r_ is the outcome metric of the reference site (either in the pre-disturbance state or a current undisturbed reference site), X_e_ is the outcome metric (e.g., abundance or diversity) after restoration (at time t = T), X_s_ is the outcome metric prior to restoration (at time t = 0) and *r* is a constant ([1/T] * Ln [X_e_/X_s_]). In the case where either X_e_ or X_s_ were zero, we replaced zero by a value in the same order of magnitude as X_s_ or X_e_ in the median magnitude (e.g., 0.5, 5, 50) (see Moreno-Mateos et al. 2017^13^). Recovery rate per annum was calculated following Jones et al., (2018)^11^ using the following equation:

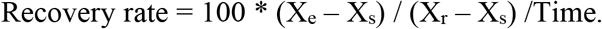

### Estimating difference between restored and unrestored habitats

To (1) estimate the difference in fish or invertebrate diversity and abundance between restored and unrestored habitats at various time-points post restoration, (2) assess differences in oyster recruitment between shell and alternate substrata, and (3) to test for the influence of relief on oyster density (by comparing adult oyster density at different reef relief), we calculated the effect size of response variables (spat density, oyster density, diversity, or abundances) by using means, standard deviations (SD), and sample sizes extracted from studies^55^. We selected to use log response ratio (lnRR) as effect size because of its capacity to detect true effects (expected value of the log-proportional change between two independent and normally distributed populations) and robustness to small sample sizes^56^. LnRR was calculated using the following equation:

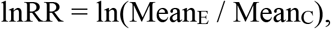

where Mean_E_ is the mean of experimental measure (e.g., number of spat on alternate substrate or adult oyster density on reef over 10 cm above sediment) and Mean_C_ treatment is the control measure (e.g. number of spat on shell or adult oyster density on reef below 10 cm on sediment). If one of the measures was zero, to avoid computational error we used a correction proportional to the reciprocal of the value of the contrasting measure (e.g: value = N, reciprocal = 1/N). When variance was reported as standard error (SE) we calculated SD as:

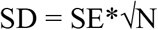

where N is the sample size. When median and ranges were reported, means and standard deviation were calculated as per Hozo et al. (2005)^57^ with the following equations:

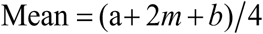

where *a* is the lower range, *b* is the upper range, and *m* is the median,

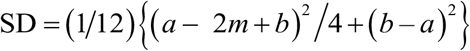

for *N* < 15, where *a* is the lower range, *b* is the upper range, and *m* is the median and

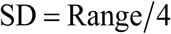

for *N* > 15. Prior to formal statistical analyses, we tested for publication bias using a Rosenberg fail-safe test, Egger’s regression test and trimfill method. Publication bias arises if studies with non-significant effects are not published^58^ and are thus excluded in analysis, thereby influencing results and interpretation. The Rosenberg fail-safe test calculates the number of studies with non-significant effects (effect size of zero) that would be required to change the results of the meta-analysis from significant to non-significant (Rosenberg 2005). The Rosenberg fail-safe numbers calculated in our analysis were larger than 5*n* + 10, where *n* is the number of studies included in the analysis^58^ and observed significance lower than 0.05 The Egger’s regression tests were used to estimate asymmetry in funnel plots and any asymmetry was adjusted using the trimfill method. For all data, either the regression tests resulted in significance values above 0.05 or the trimfill method did not change the mean effect size estimations (Supplementary File 3). Therefore, publication bias was unlikely to affect our results. Following publication bias tests, we used a weighed Random-Effects model (restricted maximum likelihood) to undertake our meta-analyses, including heterogeneity test (Q) that indicates the percentage variation between studies due to heterogeneity (i.e., differences in outcomes between different studies; also denoted as *I*^2^) rather than chance ^59^. We then performed meta-regressions using Mixed-Effects models to analyse variation in effect sizes (e.g., relationship between nekton abundance effect sizes with time post restoration). All calculation of effect sizes, publication bias tests, meta-analysis, and meta-regressions were performed on Meta-Essentials 1.5^60^ and OpenMEE, which is an open-source software specifically designed for meta-analysis in ecology and evolutionary biology and based on the “metafor” and “ape” packages for R (Wallace et al. 2017).

## Data availability

Data will be made publicly available on the University data portal (DOI will be assigned on publication).

## Acknowledgements

This project was funded by a University of Hong Kong Post-Doctoral Fellowship to DAH and an Environment and Conservation Fund Hong Kong grant (ECF106/2019), a Faculty of Science (HKU) Rising Star Fund to BDR, and an ARC Linkage Grant (LP180100732) to MJB.

