## Supplementary Figure 1 for "Global success in oyster reef restoration despite ongoing recovery debt"

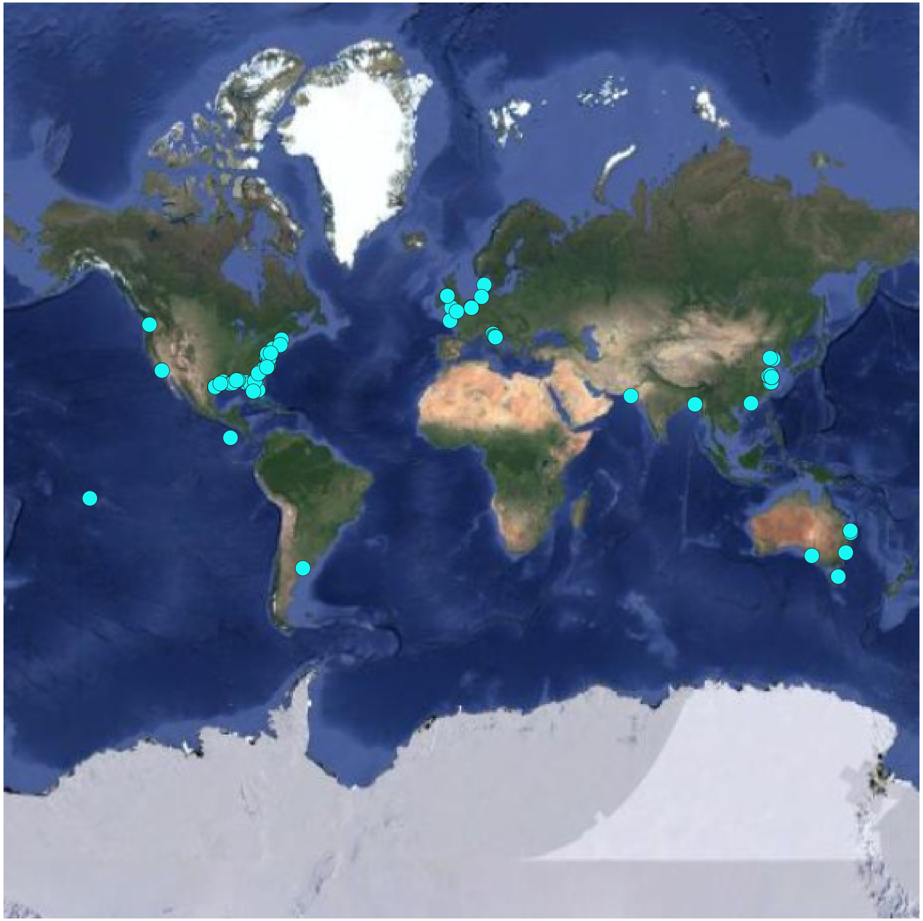

**Supplementary Figure 1**: Distribution of oyster reef restoration studies worldwide. Coordinates were extracted from 162 studies.
