## Supplementary Figure 2 for "Global success in oyster reef restoration despite ongoing recovery debt"

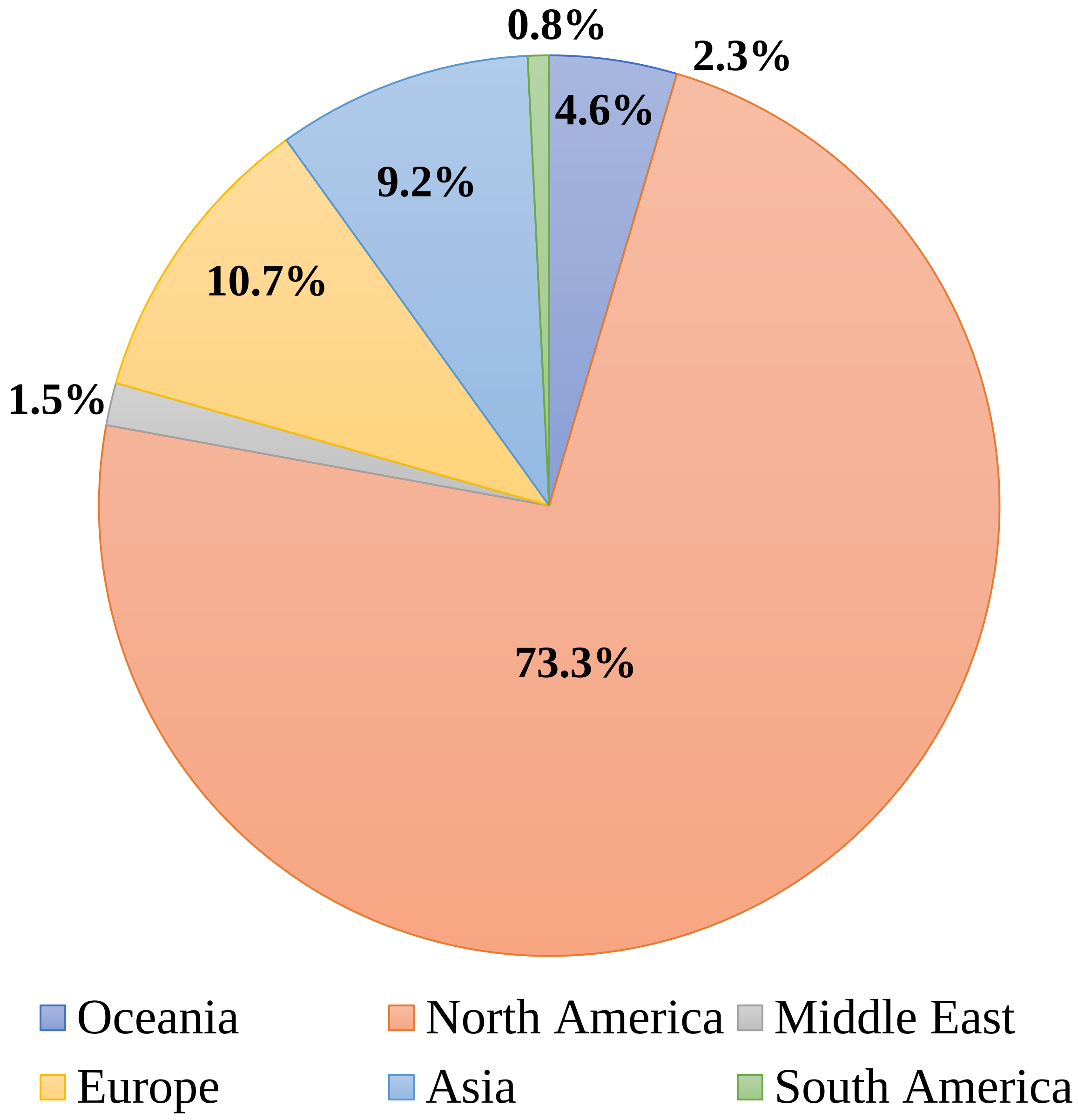


**Supplementary Figure 2:** Percentage site distribution of oyster reef projects across major areas worldwide.
